# *In-vivo* phase-dependent enhancement and suppression of brain oscillations by transcranial alternating current stimulation (tACS)

**DOI:** 10.1101/2022.02.28.482226

**Authors:** David Haslacher, Asmita Narang, Alessia Cavallo, Khaled Nasr, Emiliano Santarnecchi, Surjo R. Soekadar

## Abstract

Transcranial alternating current stimulation (tACS) can influence human perception and behavior, with recent evidence also suggesting its potential impact in clinical settings, but the underlying mechanisms are poorly understood. Behavioral and indirect physiological evidence indicates that phase-dependent constructive and destructive interference between the tACS electric field and ongoing brain oscillations may play an important role, but direct *in-vivo* validation was infeasible because stimulation artifacts impeded such assessment. Using stimulation artifact source separation (SASS), a real-time compatible artifact suppression approach, we overcame this limitation and provide direct evidence for millisecond-by-millisecond phase-dependent enhancement and suppression of ongoing brain oscillations during amplitude-modulated tACS (AM-tACS) across 29 healthy human volunteers. We found that AM-tACS enhanced and suppressed targeted brain oscillations by 11.7 ± 5.14% and 10.1 ± 4.07% respectively. Millisecond-precise modulation of oscillations predicted modulation of behavior (r = 0.65, p < 0.001). These results not only provide direct evidence for constructive and destructive interference as a key mechanism of AM-tACS but suggest superiority of phase-locked (closed-loop) AM-tACS over conventional (open-loop) AM-tACS to purposefully enhance or suppress brain oscillations.

**Significance:** The presented data provide direct evidence for a key mechanism underlying neurophysiological and behavioral effects of transcranial alternating current stimulation (tACS), a broadly used neuromodulation approach that yields promising clinical results but also raised controversies because of its variable effects. Our findings not only elucidate the underlying mechanisms of tACS, but also provide the rationale for closed-loop tACS protocols that will enable targeted enhancement and suppression of brain oscillations related to various brain functions such perception, memory or cognition. Towards this end, we introduce the technical prerequisites to establish millisecond-to-millisecond precise closed-loop tACS protocols that will be important to advance tACS as a neuroscientific and clinical tool, for example in the treatment of neuropsychiatric disorders.

## Introduction

Neural oscillations are a fundamental mechanism for precise temporal coordination between neuronal responses related to various brain functions, such as memory, perception or decision making (1, 2). Accordingly, most neuropsychiatric disorders show alterations in oscillatory brain activity that can be associated with deficits in neurocognitive function (3-6). It was shown that applying weak (1-2 mA) oscillating electric currents to the scalp in the form of transcranial alternating current stimulation (tACS) can enhance working memory capacity in the elderly (7), reduce obsessive-compulsive behavior (8), or reduce Parkinsonian tremor (9) and frequency of epileptic seizures (10). Here, the impact of tACS was largest when its frequency was tuned to the frequency of brain oscillations linked to these different brain functions or clinical symptoms. The underlying mechanisms of tACS effects are unclear, however, because stimulation artifacts impeded the assessment of millisecond-by-millisecond direct tACS effects on brain oscillations via, for instance, encephalography (EEG) recording (11-14). By assuming a direct link between behavioral outcome measures, e.g., amplitude of tremor (9), and ongoing brain oscillations, transient phase-dependent constructive and destructive interference was suggested as a key mechanism of tACS. Sophisticated study designs provided further indirect evidence for phase-dependent tACS effects on brain oscillations. For example, it was shown that tACS had a prolonged phase-specific enhancing or suppressing effect on visually evoked steady state responses (SSR) (15), and that brightness perception of flickering light depended on phase shift between sensory and electrical stimulation (16). Direct neurophysiological evidence for phase-specific constructive and destructive interference in humans, however, was not provided yet.

Here, we used an innovative approach that allows for real-time stimulation artifact rejection (17) to evaluate millisecond-by-millisecond phase-dependent tACS effects on human brain oscillations. Using electroencephalography (EEG), we assessed visually evoked SSR targeted by amplitude-modulated tACS (AM-tACS) during a binocular rivalry task (Fig. 1). In such a task, dissimilar (contradictory) visual stimuli are presented to each eye, resulting in multistable perception and fluctuating perceptual dominance of either the left or right retinal image. We asked 29 healthy human volunteers to indicate perceptual dominance of either the left or right retinal image by pressing and holding a button with either the left or right index finger. Visual stimuli presented to each eye were phase shifted by 180° and alternated in time at theta frequency (6 Hz) (18), resulting in phase-locked SSR at the same frequency. Brain oscillations were then targeted by 6-Hz AM-tACS that was phase-locked to the SSR at 0°, 90°, 180° and 270° for 20 seconds in a pseudorandom order. We hypothesized that the trial-by-trial amplitude of targeted brain oscillations (single dominance periods) would increase and decrease depending on the relative phase shift between AM-tACS and the visually evoked SSR. Moreover, we hypothesized that the modulation of trial-by-trial SSR amplitude would predict the modulation of perceptual dominance, as indicated by motor behavior.

**Figure 1.**
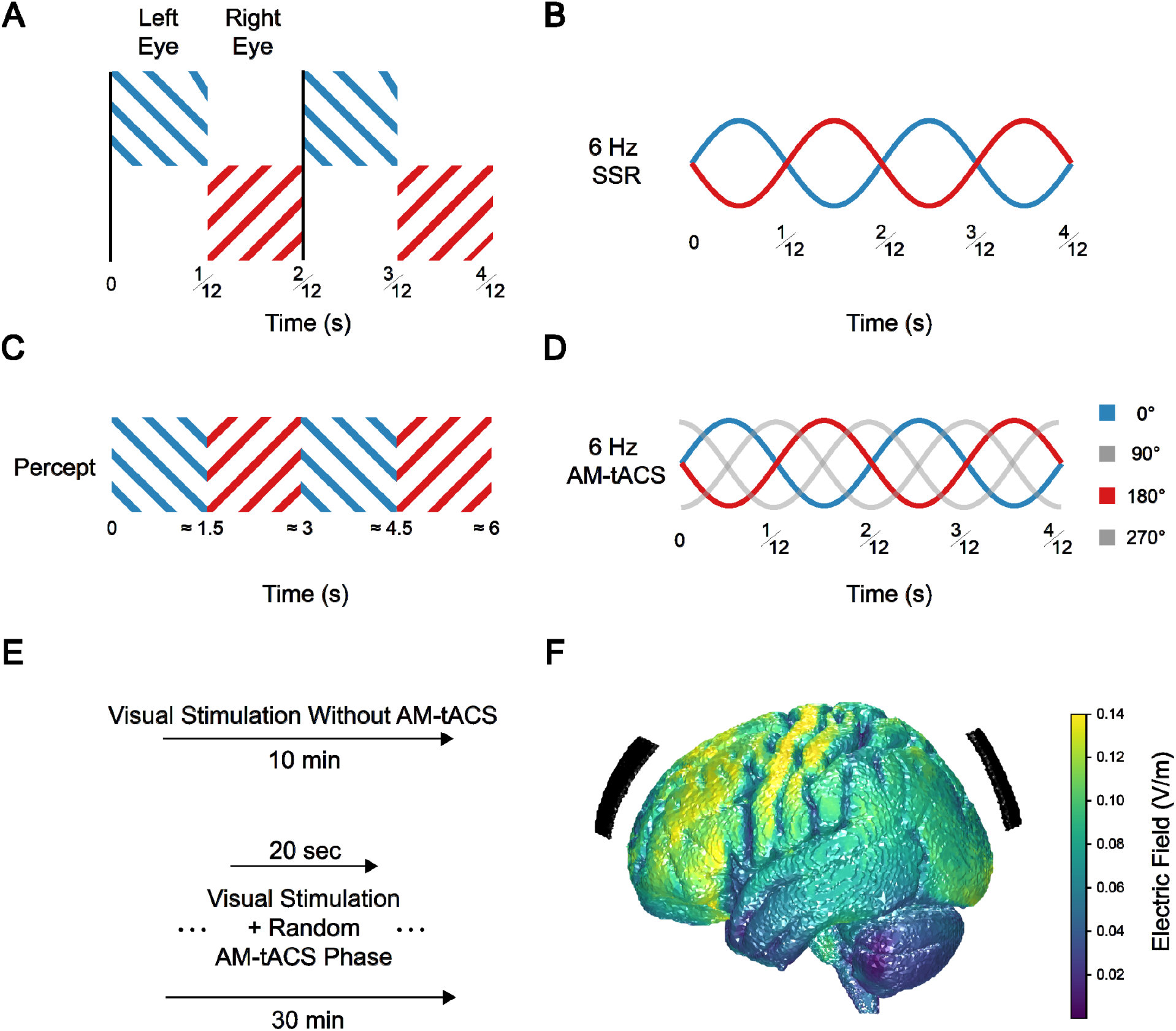
Task design and hypotheses. (A) Participants (n = 29) were presented with different visual gratings to the left (blue) and right (red) eyes. Gratings alternated in time at a rate of 6 Hz. A visual stimulus trigger (vertical black line) was used to assess selective synchronization between visually evoked steady state responses (SSR) in the electroencephalogram (EEG) and the visual stimuli. (B) We predicted that the SSR in frontal and occipital areas would synchronize with the perceptually dominant visual stimulus at opposite phases relative to the stimulus trigger. (C) During the task, conscious awareness of either the left or right visual stimulus was expected to alternate approximately every 1.5 seconds, typical for binocular rivalry. (D) Applying amplitude-modulated transcranial alternating current stimulation (AM-tACS) to the frontal and parietal lobe at different phases relative to the visual stimulus trigger was predicted to modulate the perceptual dominance ratio of the stimuli. During stimulation, fluctuation of SSR phase relative to AM-tACS was predicted to modulate SSR amplitude. (E) Prior to the application of AM-tACS, participants performed the task in absence of stimulation (for 10 minutes). During the application of AM-tACS, a new stimulation phase was chosen in a pseudorandom order every 20 seconds (for 30 minutes in total). (F) Electric field modeling showed that AM-tACS induced a broad field with highest strength in frontal and parietooccipital areas.

First, we validated that reliable single-trial reconstruction of visually evoked SSR related to left and right visual stimuli during AM-tACS was feasible. Then, we evaluated enhancement and suppression of SSR amplitude and perceptual dominance depending on the AM-tACS phase. Subsequently, we estimated the strength of constructive and destructive interference by identifying brain regions showing most prominent phase-dependent modulation of visually evoked SSR and evaluated to what extent it explained modulation of perceptual dominance. To rule out that electric stimulation of the retina could explain these modulations, we verified that subject-specific phase differences between visual stimuli and SSR predicted phase differences between AM-tACS and visual stimuli that resulted in maximal effects. We reasoned that direct retinal stimulation would not show such variable association, but rather lead to a zero- or 180°-phase relationship between AM-tACS and visual stimuli exhibiting maximal effects.

## Results

### Reconstruction of single-trial visually evoked SSR during binocular rivalry

In absence of electric stimulation, we found that, during the task, the phase opposition sum (POS), reflecting trial-by-trial synchronization of SSR with the perceptually dominant stimulus, increased in frontal and occipital brain regions across all participants (Fig. 2A). While this increase was most prominent in frontal and occipital areas, it was highly significant in most electrodes (58/64) at the group level (permutation test, p < 0.001, Bonferroni corrected). In absence of electric stimulation, a clear SSR phase opposition during left and right perceptual dominance periods was found (Fig. 2B, left panel). During AM-tACS, stimulation artifacts impeded reconstruction of the SSR, so that no such anti-phasic relationship during left and right dominance could be found (Fig. 2B, middle panel). Using SASS, an anti-phasic relationship could be successfully restored (Fig. 2B, right panel).

**Figure 2.**
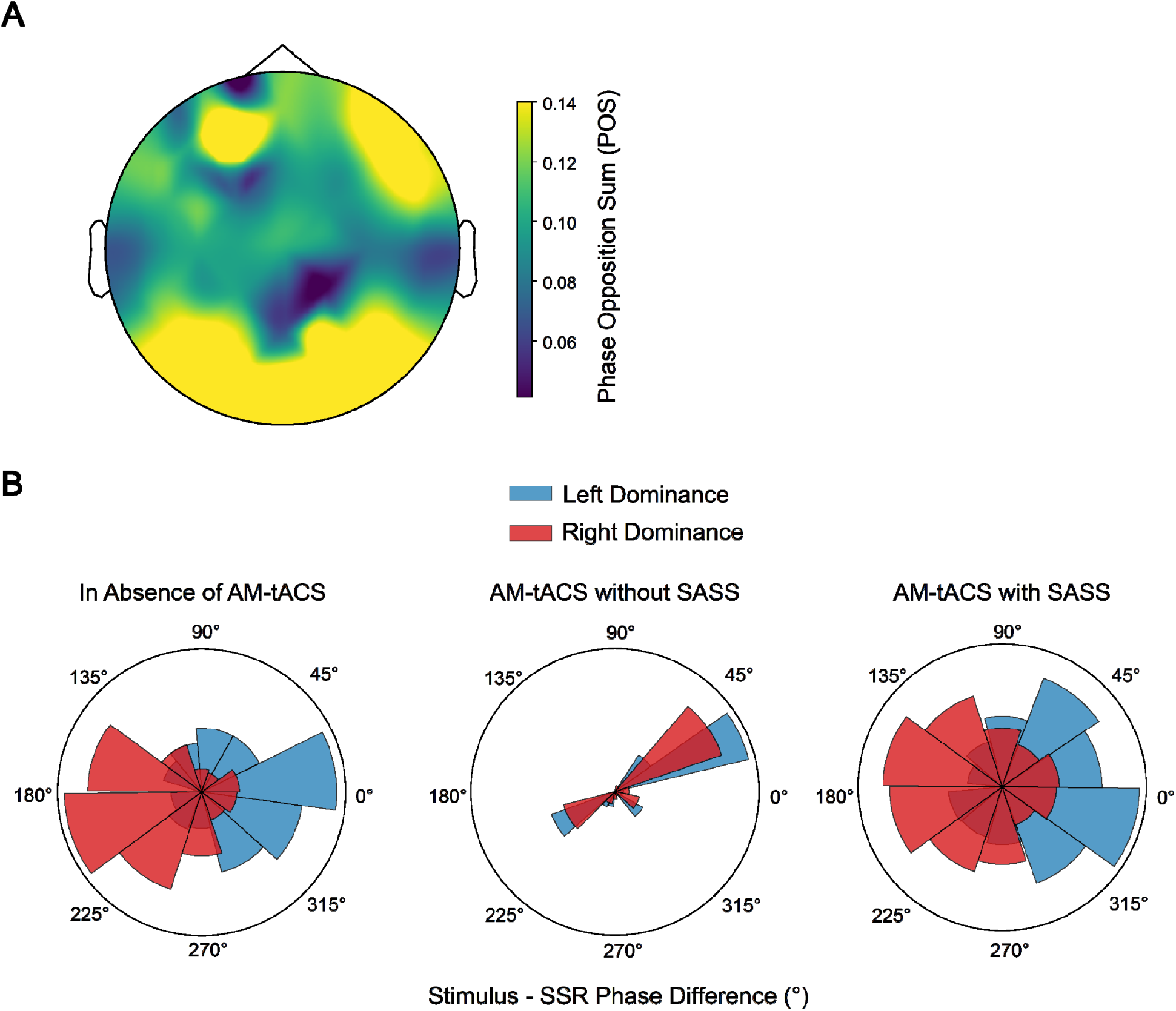
Single-trial steady-state cortical responses during binocular rivalry. (A) Steady-state responses (SSR) in frontal and parietooccipital brain areas synchronized with the perceptually dominant visual stimulus during binocular rivalry across all participants. We found that the stimulus – SSR phase difference was clustered around opposite values during left and right stimulus dominance periods. This effect was most prominent in frontal and parietooccipital areas, but was highly significant (permutation test, p < 0.0001, Bonferroni corrected) in the majority (58/64) of electrodes at the group level. (B) SASS recovered single-trial phase information in the EEG during AM-tACS (illustrated for a representative participant). In absence of AM-tACS (left panel), the stimulus – SSR phase difference was clustered around opposite values for left and right stimulus dominance periods. During the application of AM-tACS (center), the stimulation artifact obscured physiological EEG activity. Using SASS (right panel), physiological EEG activity was recovered as evidenced by the re-emergence of an anti-phasic relationship of SSR phase during left and right dominance periods.

### AM-tACS phase-dependent enhancement and suppression of brain oscillations

Evaluation of the overall impact of AM-tACS on SSR amplitudes across all participants showed an average increase of SSR amplitude by 11.7 ± 5.14% (in-phase) or suppression by 10.1 ± 4.07% (anti-phase) relative to baseline (Fig. 3A). This modulatory effect on SSR amplitude was reflected in the modulation of perceptual stimulus dominance ratio, which increased by 8.60 ± 4.90% (in-phase) or decreased by 9.06 ± 5.09% (anti-phase) (Fig. 3B).

**Figure 3.**
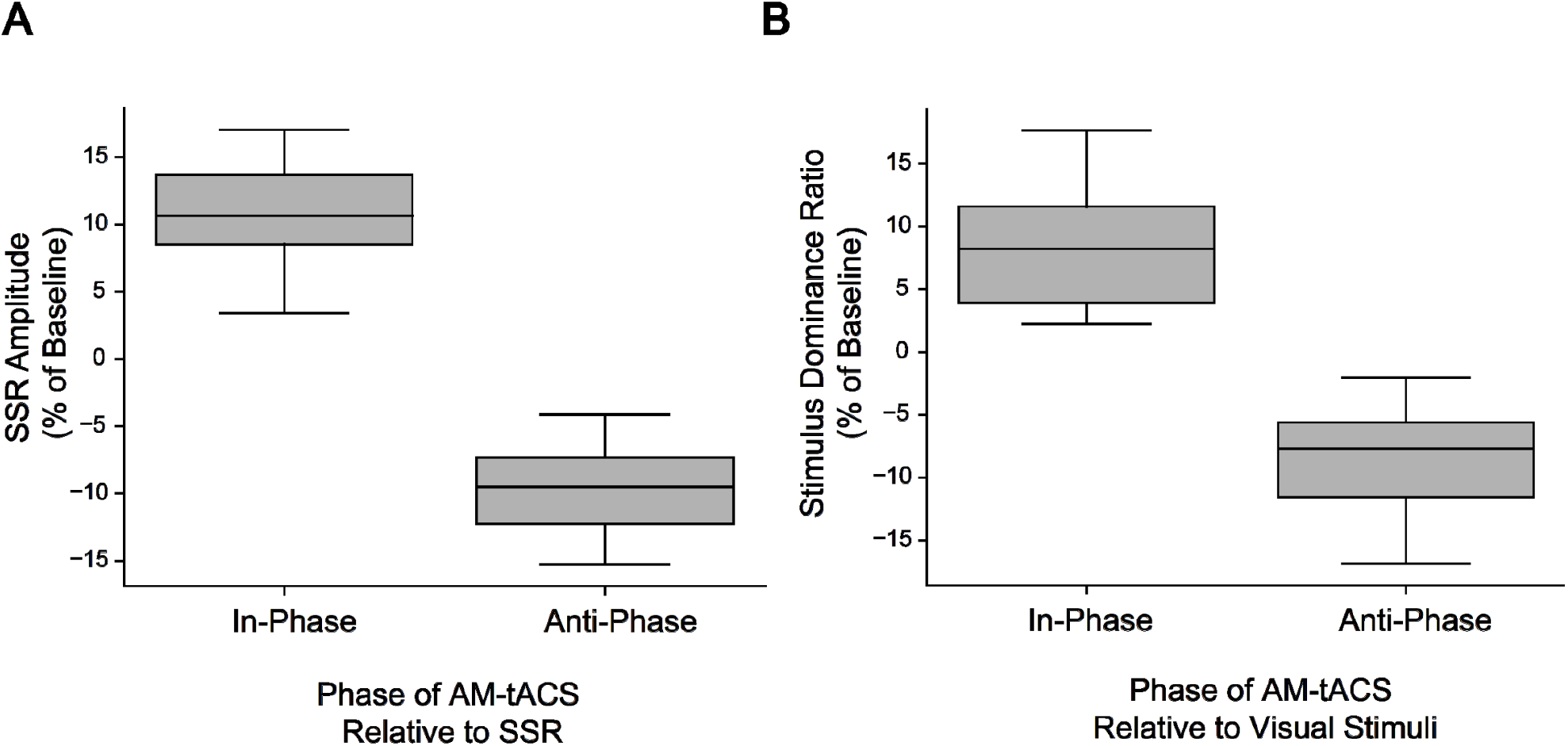
Phase-dependent enhancement and suppression of steady-state cortical responses by amplitude-modulated transcranial alternating current stimulation (AM-tACS). (A) When AM-tACS was applied at an optimal phase angle relative to SSR (in-phase), SSR amplitude was enhanced by 11.7 ± 5.14 % relative to baseline. When AM-tACS was applied at the opposite phase angle (anti-phase), amplitude of SSR was suppressed by 10.1 ± 4.07 %. (B) When AM-tACS was applied in-phase relative to the visual stimulus trigger, the perceptual dominance ratio was enhanced by 8.60 ± 4.90 %, while it was suppressed by 9.06 ± 5.09 % when AM-tACS was applied anti-phase.

Evaluating single-trial SSR amplitudes showed a clear increase or decrease depending on the phase difference between AM-tACS and SSR (permutation test, p < 0.01) (Fig. 4A).

**Figure 4.**
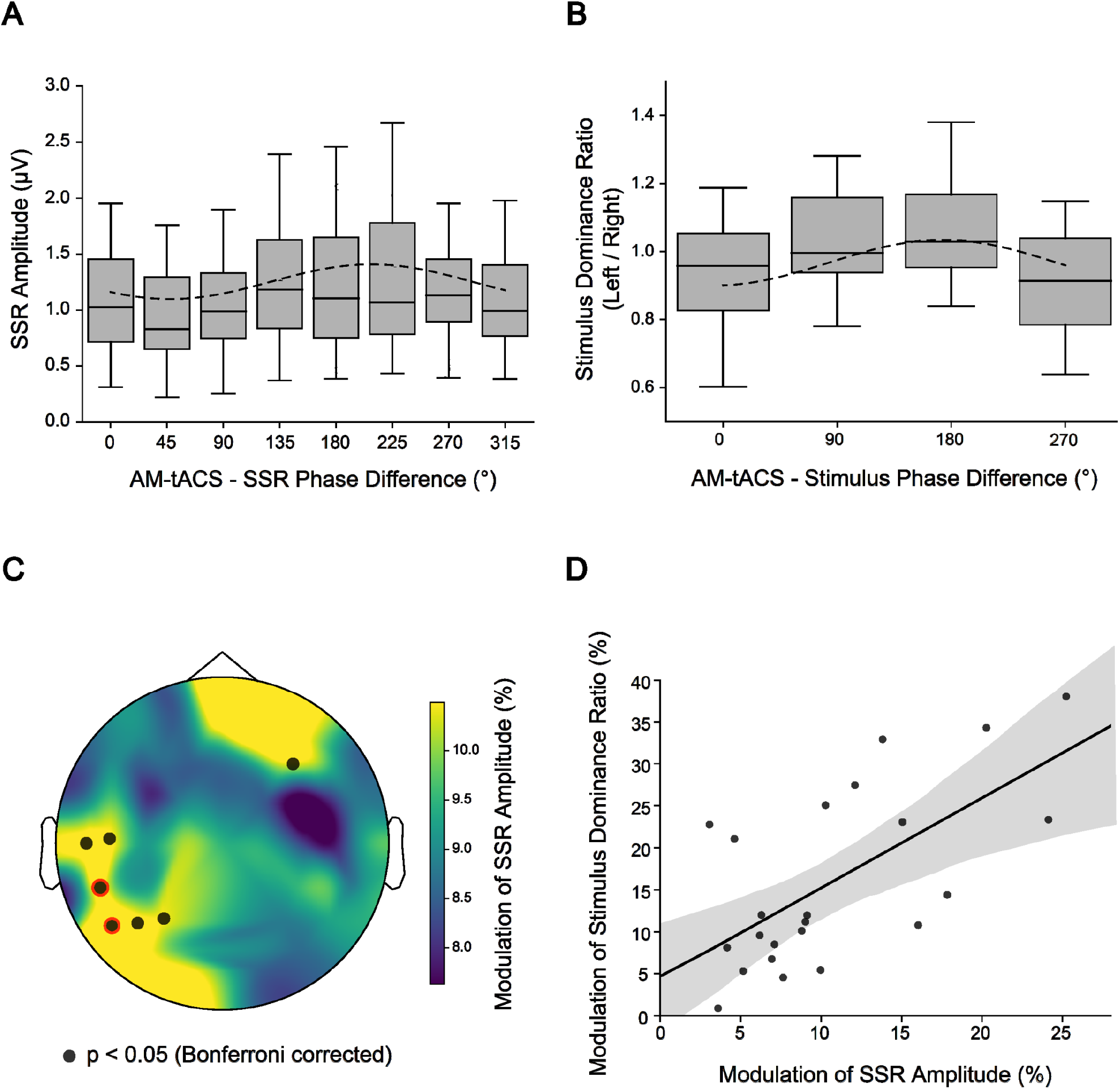
Electric stimulation effects on single-trial SSR amplitude and perception. (A) Amplitude-modulated transcranial alternating current (AM-tACS) modulated SSR amplitude in a selected participant during single dominance periods in a phase-dependent manner (permutation test, p < 0.01). (B) AM-tACS modulated the perceptual dominance ratio of visual stimuli in a phase-dependent manner (permutation test, p < 0.05). As the optimal stimulus-tACS phase difference varied across participants (Fig. 5A), phase bins were aligned across participants for visualization. (C) Phase-dependent modulation of SSR amplitude was found in frontal and temporo-parieto-occipital areas (permutation test, p < 0.05, Bonferroni corrected), which predicted the perceptual modulation (Pearson’s r, p < 0.05, Bonferroni corrected). Electrodes marked in black indicate significant modulation of SSR amplitude, while electrodes marked with a red circle indicate significant modulation of theta amplitude in addition to significant correlation with modulation of perception. (D) Phase-dependent modulation of SSR amplitude predicted the phase-dependent modulation of perception (r = 0.65, p < 0.001) in one of the electrodes marked with a red circle (electrode position TP7 according to the international 10/20 system), explaining 42% of the inter-individual variance in perception.

In accordance with this effect, AM-tACS showed an impact on the perceptual dominance ratio depending on the phase difference to visual stimuli across all participants (permutation test, p < 0.05) (Fig. 4B).

A relationship between modulation of SSR amplitude and modulation of perceptual dominance ratio was found across all participants and was most prominent in the left temporo-parieto-occipital region (Pearson’s r = 0.65, p < 0.001, Bonferroni corrected) (Fig. 4C and 4D). While the phase-dependent modulation depth of SSR amplitude varied across participants (11.0 ± 6.05%) at this location (electrode position TP7 according to the international 10/20 system), the modulation of SSR amplitude explained 42% of the inter-individual variance in the modulation of the perceptual dominance ratio.

### Subject-specific phase differences between visual stimuli and cortical responses

By calculating the circular correlation between the visual stimulus - SSR phase difference in absence of electric stimulation with the AM-tACS – visual stimulus phase difference that resulted in maximal behavioral effects across all participants, we found that this correlation was highest in left parietooccipital electrodes (O1 and PO7 according to the international 10/20 system) (cluster-based permutation test, rho = 0.52, p < 0.05) (Fig. 5A).

**Figure 5.**
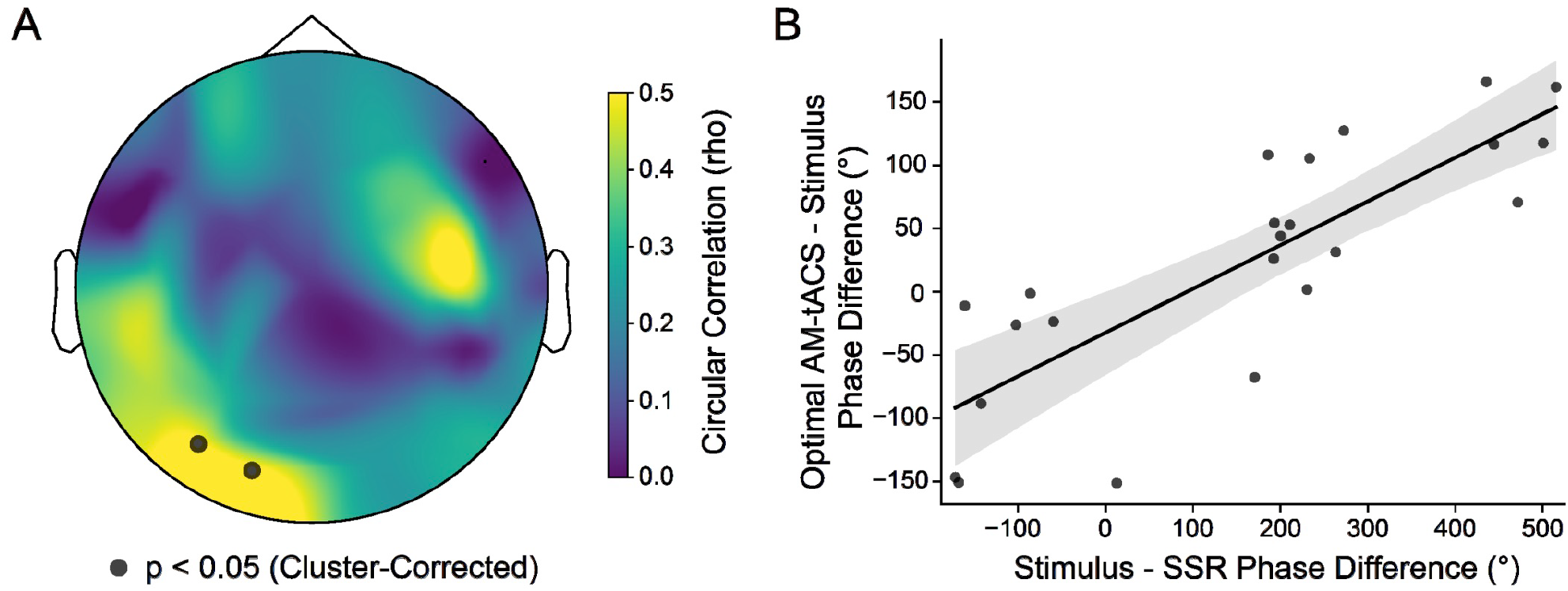
Subject-specific phase differences between visual stimuli and cortical responses. (A) Parieto-occipital electrodes (electrode positions O1 and PO7 according to the international 10/20 system indicated as black dots) showed significant circular correlation between the visual stimulus - SSR phase difference in absence of electric stimulation and the AM-tACS – visual stimulus phase difference that resulted in maximal behavioral effects (cluster-based permutation test, rho = 0.52, p < 0.05). (B) For one of these electrodes (O1), this relationship is visualized. The optimal AM-tACS – visual stimulus phase differences varied across participants, indicating that modulation of cortical responses and not direct retinal or sensory stimulation caused a substantial portion of the reported neurophysiological and behavioral effects.

We found that the optimal phase difference between AM-tACS and visual stimulus varied across participants and was not zero or 180° (Fig. 5B) indicating that direct modulation of cortical responses and not retinal activation caused the reported neurophysiological and behavioral effects.

## Discussion

Our results show that physiological and behavioral effects of AM-tACS depend on constructive and destructive interference between the electric field and targeted brain oscillations. To enable highly specific targeting in terms of phase, we used visually evoked cortical responses in combination with AM-tACS. By transiently shifting the phase between AM-tACS and visual stimuli in a random order, we could – using SASS – directly assess the impact of such a phase shift on brain physiology and behavior. Besides elucidating the underlying mechanisms of AM-tACS, our study exemplifies the capability of concurrent tACS-EEG for investigating brain-behavior-relationships. Since our approach is real-time compatible, it paves the way for closed-loop or adaptive transcranial electric stimulation paradigms that selectively enhance or suppress targeted brain oscillations. While enhancing brain oscillations, e.g., in the alpha band, has shown to mediate default mode network (DMN) connectivity enhancement (19), or, in the case of frontal theta oscillations, modulate working memory and cognitive function (20, 21), suppression of brain oscillations has been mainly described in the context of deep brain stimulation (DBS) ameliorating clinical symptoms, e.g. in Parkinson’s or dystonia (22, 23). Our results suggest that closed-loop or adaptive AM-tACS to suppress targeted oscillations might open a new avenue to ameliorate such symptoms non-invasively. This may also play a role for targeting pathological brain oscillations found across various neuropsychiatric disorders, such as depression (24), obsessive compulsive disorder (OCD) (25), attention deficit and hyperactivity disorder (ADHD) (26, 27), or Alzheimer’s disease (28).

Since our study was specifically designed to investigate phase-dependent effects of AM-tACS on steady-state brain responses, the presented results do not allow to draw any conclusions about other possible mechanisms of immediate or delayed tACS effects, such as entrainment (29), resonance (30) or plasticity in oscillatory circuits (31). Our results underline, however, that the interaction between tACS and ongoing brain activity could involve antagonizing effects, such as transient enhancement and suppression of oscillatory activity, that might explain variability of tACS effects reported for various experimental paradigms where phase information was not taken into account (32, 33).

Retinal or sensory stimulation have been identified as a common confound in tACS studies (34). For instance, it was shown that electric currents can spread over the scalp and directly stimulate the retina (35). Translated to our paradigm, simultaneous (zero-phase difference) visual and electric stimulation of the retina would result in largest cortical responses and brightness perception, while anti-phasic (180° phase difference) visual and electric stimulation would result in attenuated responses and brightness perception. Besides direct stimulation of the retina, also somatosensory stimulation via peripheral nerves can alter brain activity during tACS (36). In analogy to direct retinal stimulation, it was found that rhythmic somatosensory stimulation optimally enhances perceptual dominance during binocular rivalry at zero-phase difference between visual stimuli and sensory stimulation (37). This was similarly shown in multisensory integration across the auditory and visual domain (38).

We found, however, that the phase difference between AM-tACS and visual stimuli leading to maximal modulation of the perceptual dominance ratio varied across individuals and could be predicted across participants by the latency between visual stimuli and SSR in absence of AM-tACS. This corroborates direct modulation of cortical responses and excludes that retinal or sensory stimulation caused the described effects.

An important limitation of our study relates to the generalizability of our findings to other tACS protocols, particularly monosinusoidal tACS. While it is very likely that also monosinusoidal tACS results in phase-dependent enhancement and suppression of brain oscillations at the target frequency, lack of a reliable strategy to recover phase and amplitude during such stimulation protocol impedes further validation. While other AM-tACS protocols used higher carrier frequencies (e.g., 220 Hz) (39), a 40-Hz carrier frequency was used to mimic the ubiquitous theta-gamma cross-frequency coupling in the human neocortex (40, 41). It is unclear whether other carrier frequencies would have led to other results, an issue that should be investigated in future studies. Another important question relates to the dose-response relationship between transcranial electric stimulation intensity and physiological as well as behavioral effects. The strategy used in this work allows now to systematically investigate this controversial issue (42).

## Materials and Methods

### Participants

This study was performed with healthy human volunteers at the Charité – Universitätsmedizin Berlin, Germany. In total, 29 participants (13 female, 16 male) between the age of 21 and 36 (mean age 26 ± 4 years) were recruited. All participants provided written informed consent in accordance with the ethics committee of the Charité – Universitätsmedizin Berlin (EA1/077/18). Participants were compensated with 10 Euros per hour. Data from two participants had to be excluded because of technical defects during the experiment (cable breakage, software crash). Data of two participants were excluded due to a short-circuit between the stimulator and most EEG electrodes (probably due to extensive sweating).

### Electroencephalography (EEG)

EEG was recorded throughout four 10-minute experimental blocks using a 64-channel system (Bittium Corp., Oulu, Finland) with electrodes positioned according to the international 10-20 system. For all recordings, the amplifier was set to DC-mode with a dynamic range of +/-430 V, a resolution of 51 nV/bit, and a range of 24 bit. EEG was recorded at 500 Hz with an anti-aliasing filter applied at 125 Hz. Electrode impedances were maintained below 10 kOhm or excluded from the analysis.

### Preprocessing of EEG data

EEG analyses were performed using the MNE-Python software (43), an open-source Python package for analyzing human neurophysiological data. Recordings were re-sampled at 100 Hz for further processing. All EEG recordings were initially bandpass-filtered from 5-7 Hz with a finite impulse response (FIR) filter designed using the Hamming window method (44). For data recorded in the presence of AM-tACS, stimulation artifact source separation (SASS) was applied to remove the stimulation artifact (45). SASS was applied in non-overlapping windows of 30 seconds, for each of which the artifact-contaminated covariance matrix A and SASS projection matrix P were recomputed. The covariance matrix B of data recorded in absence of AM-tACS remained constant. To select the number of components to reject, |*corr*(*B*) − *corr*(*PAP*^*T*^)|_2_ was minimized, where corr refers to the conversion of a covariance matrix into a correlation matrix, and *PAP*^*T*^ represents the covariance matrix of data after application of SASS. Data was then re-referenced to the common average reference. To obtain instantaneous phase and amplitude, the Hilbert transform was applied to the EEG data and demeaned electric stimulation envelope signal.

### Phase locking of visual stimuli and visually evoked steady-state responses (SSR)

To assess phase-locking between visual stimuli and SSR, the instantaneous phase of EEG signals at the timepoints of the visual stimulus trigger were computed in a 500 ms period centered to the onset of perceptual dominance periods of the left and right visual stimuli. The circular mean was used to average phase angles within each perceptual dominance period. The phase opposition sum (POS) (46) was then used to contrast SSR phase during left and right perceptual dominance periods.

### Perceptual dominance ratio

For all AM-tACS stimulation conditions, a perceptual dominance ratio was obtained. This was calculated as 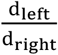, where d is the total time of perceptual dominance indicated by pressing and holding a button with either the left or right index finger.

### Impact of AM-tACS on SSR and perceptual dominance

To assess modulation depth of visually evoked SSR, instantaneous SSR amplitude was averaged within each dominance period of left and right visual stimuli. Likewise, instantaneous phase differences between AM-tACS and SSRs were averaged within each dominance period using the circular mean. Then, dominance periods were assigned to eight equidistant phase bins. Consequently, the mean SSR amplitude within each phase bin was computed, and a discrete Fourier transform was used to obtain the modulation amplitude, which was converted into a percent of the mean SSR amplitude across phase bins. To assess the AM-tACS-caused modulation depth of the perceptual dominance ratio, a discrete Fourier transform was used to obtain the modulation amplitude, which was converted into a percent of the mean perceptual dominance ratio across phase bins. To compute maximal (in-phase) and minimal (anti-phase) SSR (Fig. 3A) and perceptual (Fig. 3B) modulation, the percent change of the average within the maximal and minimal phase bins were computed relative to the average across all phase bins.

### Statistical analysis

To assess significance of the phase opposition sum (POS), a permutation test was applied. POS was computed for 1000 random permutations of “left” and “right” labels over dominance periods, from which a p-value for the true POS value was obtained. To obtain a group-level p-value for the aforementioned statistics at each EEG electrode, Fisher’s method was employed to combine participant-level p-values (47). To test for AM-tACS-caused sinusoidal modulation of SSR amplitude, we employed permutation statistics (48). For each participant and EEG electrode, a discrete Fourier transform was used to obtain the true modulation amplitude. Then, 1000 random permutations of the data across phase bins were employed to obtain a surrogate distribution of modulation amplitudes, from which a p-value was obtained. To combine p-values across participants, Fisher’s method was used. A permutation test was also used to test for AM-tACS-caused modulation of the perceptual dominance ratio. A discrete Fourier transform was used to obtain the true modulation amplitude of perceptual dominance ratio. Then, 1000 random permutations of the data across phase bins were employed to obtain a surrogate distribution of modulation amplitudes, from which a p-value was obtained. To assess the correlation across participants between the SSR and perceptual modulation depth at each EEG electrode, Pearson’s r was used. For all statistical tests performed individually for each EEG electrode, Bonferroni correction was applied to account for multiple comparisons across electrodes. To plot the modulation of perceptual dominance ratio (Fig. 4B), phase bins for the participant-specific sine fits were aligned for visualization purposes after performing statistical hypothesis testing. While the phase of AM-tACS generally modulated SSR and perceptual outcome measures on a single-participant level, the optimal phase for this modulation can differ across participants (48). In each iteration of the cluster-based test, a t-test was performed for each electrode, comparing the distribution (across participants) of average SSR amplitude during AM-tACS. The t-statistic corresponding to an alpha level of 0.05 was used to threshold each electrode and obtain contiguous clusters. The cluster-level statistic then consisted of the average t-statistic in each cluster. This procedure was then repeated 1000 times with a permutation of SSR amplitudes between conditions to obtain a null distribution of cluster-level statistics from which p-values for each cluster were then obtained by comparison with the true cluster-level statistic. A similar procedure was employed to test for circular correlation between stimulus-locked SSR phase and optimal AM-tACS phase (Fig. 5B). Here, however, circular-circular correlations were tested at each electrode (49), and permutations were performed between participants.

### Visual stimulation and perceptual response

Participants were presented with a green grating rotated 45° counterclockwise from a vertical position displayed to their left eye, and a red grating rotated 45° clockwise displayed to their right eye. Stimuli were displayed separately to each eye using a head-mounted display (Oculus VR Inc., California, USA). Left and right gratings were presented alternatingly at 6 Hz on a black background, with a red fixation dot in the center. Consistent with a refresh rate of 60 Hz, each stimulus was therefore displayed for three frames while the other stimulus was absent. A visual stimulus trigger was recorded in the EEG system that was synchronized with the onset of the left visual stimulus. Participants experienced binocular rivalry, i.e., perceptual dominance of either the left or the right grating. They were asked to indicate dominance of the left or right grating by pressing and holding a button with either the left or right index finger. Participants were asked to refrain from pressing any keys during phases of mixed perceptual dominance.

### Electrical stimulation

Amplitude-modulated transcranial alternating current stimulation (AM-tACS) was applied over the frontal and parietal lobe with a 6 Hz envelope, 40 Hz carrier frequency, and peak-to-peak amplitude of 2 mA. Rectangular rubber electrodes with dimensions 5 × 7 cm were placed perpendicular to the midline, centered on positions AFz and Pz of the international 10-20 system, and applied using conductive ten20 paste (Weaver & Co, Aurora, CO, USA). AM-tACS was applied at four different equidistant phase angles (0°, 90°, 180°, 270°) relative to the visual stimulus trigger. Every 20 seconds, a new AM-tACS phase was chosen in a pseudorandom manner, ensuring that each phase was applied for an equal duration. AM-tACS was applied using a DC-STIMULATOR PLUS (NeuroConn GmbH, Ilmenau, Germany) that outputs a current proportional to a provided voltage. The AM-tACS waveform was generated by a Simulink Real-Time target machine connected to a 16-bit analog input/output module (Speedgoat GmbH, Liebefeld, Switzerland). The real-time target machine received an analog trigger signal from the head-mounted display to ensure correct synchronization with the visual stimuli at the desired phase angle.

### Recording blocks

Participants participated in four consecutive recording blocks of 10 minutes each, during each of which they performed the binocular rivalry task. The first session was performed in absence of AM-tACS, whereas the last three sessions were performed while AM-tACS was delivered.

## Acknowledgements

This research was supported in part by the European Research Council (ERC) under the project NGBMI (759370), ERA-NET Neuron under the project HYBRIDMIND (01GP2121B) and the Einstein Stiftung Berlin.

